# Ecogenomics of transcontinental black spruce: identification of climate adaptation genes across the Canadian boreal landscape

**DOI:** 10.64898/2026.03.26.714629

**Authors:** Vincent Quevillon, Sébastien Gérardi, Patrick Lenz, Jean Bousquet

## Abstract

Black spruce (*Picea mariana* [Mill.] B.S.P.) is an emblematic and ubiquitous species of the North America’s boreal forest. While conifer breeding programs have traditionally focused on growth and wood property traits, the study of climate adaptation traits is becoming increasingly prevalent, given the predicted impact of climate change on North America’s boreal zone. Through this study, we aimed to identify genes associated with climate adaptation in black spruce across Canada. A total of 254 black spruce trees from 30 populations, covering most of the species’ distribution range, were sampled and genotyped for SNPs located in ∼5000 gene loci. Uni- and multivariate Genotype-Environment Association (GEA) approaches, namely LFMM and RDA, as well as an outlier method based on population differentiation (*F*_ST_) were used to identify genes significantly associated with climatic factors. As such, a total of 77 genes carrying significant candidate SNPs were identified, among which 14 candidates were corroborated by at least two methods. Many of these gene SNPs were also confirmed at a smaller geographic scale, across west - east partitions corresponding to the two main black spruce historical lineages. Notably, significant gene SNPs were more frequently associated to moisture/aridity factors in the western part of the range, and more to temperature factors in the eastern part. The genes carrying these SNPs were also frequently associated to abiotic and biotic stress response. In the context of rapid climate change in the Canadian boreal forest, the results obtained within the framework of this study should support implementing gene conservation efforts while assisting prediction in black spruce breeding programs, which are instrumental to producing adapted planting stock for the large-scale reforestation efforts conducted annually across the Canadian boreal forest.

## Introduction

Predicted climate change in all current plausible scenarios poses significant risk to forest ecosystems on a global scale (Intergovernmental Panel on Climate Change - IPCC, 2023). In North America’s boreal forest, this could translate into an increase of the frequency and severity of abiotic disturbances, including ice storms, droughts, permafrost thaw and wildfires (Williamson et al., 2009; Price et al., 2013; Francis & Vavrus, 2015; Cook et al., 2018; Boucher et al., 2018), as well as biotic disturbances such as pest insect outbreaks, spread of fungal-related diseases and invasive species introduction (Lemprière et al., 2008; Williamson et al., 2009). These outcomes could lead to substantial ecological and economic repercussions given the importance of North America’s boreal forest in both these regards (Wells et al., 2020). Considering this, and given the long generation time of boreal conifers (Bouillé & Bousquet, 2005), understanding the molecular basis of climate adaptation becomes increasingly valuable, particularly in the contexts of genetic conservation and tree breeding, where improving predictions of adaptive capacity under future climates becomes a priority.

Using phenotypic data gathered from common gardens, studying the association between phenotypic traits and local environmental conditions of geographic origins of candidate populations and genetic families has long been the gold standard to detect patterns of genetic adaptation to climate in a context of long-term testing in tree improvement programs (Depardieu et al., 2020). Advances in genomic research now enable to decipher the molecular basis of such genetic adaptation (Hoban et al., 2016; Depardieu et al., 2021). Given newly developed and more affordable sequencing and genotyping technologies, various genomic approaches can now be applied to non-domesticated species with large and complex genomes such as conifers (Bousquet et al., 2021). Among these, Genome-Wide Association Studies (GWAS) are becoming commonly to uncover adaptation at the molecular level in such organisms, including GPA (Genotyped-Phenotype Association) studies aim to identify relationships between genotypes and phenotypes (Bousquet et al., 2021; Uffelmann et al., 2021). Such studies have been successfully conducted on a range of North American conifers, including white spruce (*Picea glauca*) (e.g. Depardieu et al., 2021), sugar pine (*Pinus lambertiana*) (e.g. Eckert et al., 2015), eastern white pine (*Pinus strobus*) (e.g. Housset et al., 2018), or loblolly pine (*Pinus taeda*) (e.g. Talbot et al., 2017), resulting in highly useful information for tree breeding and genetic conservation efforts. Yet, GPA approaches require that individuals are grown in homogeneous environmental conditions such as in common gardens, in order to assess the contribution of genotypic information to the total phenotypic variance, which makes it impractical when natural populations or species are not represented in common garden studies, especially when these species are long-lived woody perennials. Phenotyping over many years in such common gardens also remains very costly and labor intensive.

Identifying molecular drivers of genetic adaptation without relying on phenotypic trait data still remains a challenge in evolutionary genomics and plant breeding. When phenotypes are not available, one of the most frequently used strategies to identify genetic variations putatively linked to environmental adaptation implicates methods based on the population differentiation index (*F*_ST_) of molecular marker allele frequencies (Bousquet et al., 2021). The aim of such analyses is to detect genetic loci with outlier *F*_ST_ values among a distribution of mainly neutral loci (Namroud et al., 2008; Nosil et al., 2009). Statistically high differentiation at distinct loci is then interpreted as indicative of divergent natural selection given that this mechanism can operate at distinct loci contrary to genetic drift, migration or changes in the mating system, which affect the whole genome (Lewontin & Krakauer, 1973). Thus, neutral loci should follow a general pattern dictated by neutral forces such as drift-migration equilibrium and isolation-by-distance, while loci under significant natural selection pressure should deviate significantly from these patterns (Lewontin & Krakauer, 1973; for a review, see Chung et al., 2023). This approach has been refined by Excoffier et al. (2009) to better account for historical population structure and isolation-by-distance (IBD), thus limiting the detection of false positives. Previous studies in spruce species (*Picea* sp.) relied on this type of analysis to identify gene markers related to environmental clines or conditions (e.g. Namroud et al., 2008; Prunier et al., 2011; for a review, see Bousquet et al., 2021).

Similarly, the use of landscape genomics methods, such as Genotype-Environment Association studies (GEA), has become increasingly used in the past decade. These approaches rely on the detection of statistically significant associations between allele frequency and variation in environmental factors to determine if a specific genetic variant confers a selective advantage in certain local environments (Lasky et al., 2023). However, the effect of population structure among sampled individuals must be controlled to avoid confounding effects unrelated to natural selection. Since they use both genetic and environmental variables in their underlying models, these approaches establish a more direct link between genetic variants and causal environmental factors compared to *F*_ST_-based methods. Such approaches have also been applied successfully to conifer species such as loblolly pine (e.g. Eckert et al., 2010), as well as in several spruce species (e.g. Namroud et al., 2008; Prunier et al., 2011, 2012; Chen et al., 2012; Hornoy et al., 2015; Depardieu et al., 2021; for a review, see Bousquet et al., 2021). Furthermore, given the rapid decay of linkage disequilibrium (LD) in conifer natural populations (Pavy et al., 2012), which are typically characterized by high gene flow and large population sizes (such as in boreal spruces, see Bouillé & Bousquet, 2005), and considering the predominance of non-coding DNA in conifer giga-genomes (De La Torre et al., 2014), focusing on gene-based SNPs in GEA (and GPA) approaches should enable a more precise identification of candidate genes underlying significant associations arising from cis- or trans-acting genetic effects (Prunier et al., 2012, 2015).

Black spruce (*Picea mariana*) is a largely non-domesticated and key tree species of the North American boreal forest (Brandt, 2009; Lamhamedi & Bernier, 1994). Along with other spruce species, black spruce is a major component of reforestation efforts across Canada (National Forestry Database, 2023) and it is the most widely planted tree species, with tens of millions of seedlings reforested annually in some jurisdictions, such as in the province of Quebec (Ministère des Ressources naturelles et des Forêts, 2022).

At the phylogenetical level, black spruce belongs to the North American *Picea* clade, but it forms a distinct subgroup with red spruce (*Picea rubens*) with which it hybridizes in the southeastern part of its range (Perron & Bousquet, 1997; de Lafontaine et al., 2015). This subgroup is distinct from the clade that includes white spruce and its closely related species, from which black spruce is reproductively isolated (Bouillé et al., 2011). It is found in a wide array of habitats across its transcontinental distribution, ranging from xeric sites to wetlands (Lamhamedi & Bernier, 1994), and can even persist in the harsh environmental conditions of the Arctic tundra under extreme environmental stress (Laberge et al., 2000). It is also characterized by large population sizes promoted by wind pollination, high gene flow and an outcrossing mating system (Perry & Bousquet, 2001; Bouillé & Bousquet, 2005), which translates into lowl population differentiation and high levels of heterozygosity (Isabel et al., 1995; Gamache et al., 2003; Bousquet et al., 2021). Black spruce ability to withstand a variety of environmental conditions and its high genetic diversity make it an ideal candidate for reforestation initiatives and genetic improvement programs (Mullin et al., 2011). Furthermore, recent studies on black spruce growth under various climate change scenarios showed significant variations in population responses, suggesting that intraspecific genetic diversity plays a key role in this species’ ability to cope with changing environmental conditions (Robert et al., 2024, 2025).

Black spruce possesses a giga-genome of 18.7 billion base pairs (Lo et al., 2024), and its exome (Pavy et al., 2016) and transcriptome (Pavy et al., 2025) have been well characterized. Along with white spruce, black spruce exhibits one of the highest levels of molecular genetic diversity (as per gene SNPs) among conifers (Pavy et al., 2025). Using limited SNP datasets for outlier and GEA analyses, signatures of local selection postdating Holocene recolonization were detected in a range of black spruce genes (Prunier et al., 2011, 2012). However, the overall molecular basis of climate adaptation remains poorly understood in this species as in most other conifers. Thus, studying the genomic basis of environmental adaptation using large gene sets across most of the transcontinental range of black spruce could guide gene conservation strategies and support tree breeding and management efforts aimed at climate change mitigation, maintaining ecosystem resilience, and supporting the reforestation of locally well-adapted stocks (Prunier et al., 2011; Lemprière et al., 2013; Högberg et al., 2021).

The objective of this study was to identify key candidate genes putatively involved in local adaptation in black spruce. Previous phylogeographic studies of this largely distributed species based on organelle genomes indicated that the species is divided in a few historical genetic lineages shaped by vicariance during glacial cycles (Jaramillo-Correa et al., 2004; Gérardi et al., 2010). By sampling SNPs from ∼5000 distinct nuclear gene loci, we first assessed population structure to confirm the presence of the previously reported two main historical lineages along the west-east transect of the Canadian boreal forest. Taking this structure into account, we then identified gene polymorphisms linked to climate adaptation across most of the species’ range by using GEA and *F*_ST_-based outlier detection methods. Given the different drivers of climatic variation across the Canadian boreal forest landscape, we also investigated if climatic variables implicated in local adaptation differed between regions. Functional annotations of the identified candidate genes for climate adaptation were further examined to evaluate their potential biological significance.

## Materials and Methods

### Population sampling, climatic data, and sample genotyping

#### Population sampling

Using material from the study of Prunier et al. (2012), 309 individuals from 35 black spruce natural populations, covering most of the species’ boreal range and major historical lineages, were genotyped. Samples from the suture zone between the major Eastern and Western historical lineages (provinces of Saskatchewan and Manitoba; as determined by Jaramillo-Correa et al., (2004), Gérardi et al. (2010), and Prunier et al. (2012) were deliberately excluded to avoid possible confounding effects resulting from lineage admixture. We also did not include the Maritime provinces in eastern Canada, where black spruce and red spruce co-occur and naturally hybridize (Perron & Bousquet, 1997; de Lafontaine et al., 2015). Information on sampled populations can be found in Table S1.

#### Population climatic data

Climatic data for sampled populations (Table 1) was gathered using BioSIM11 (Régnière et al., 2017). This software interpolates climatic data from the closest weather stations to estimate actual on-site conditions. Data from a 30-year period from 1950 to 1980 was collected. This corresponds to the period when mother trees from which seed collections were originally made reached their sexual maturity. In total, 26 climatic factors were extracted (Table 1). These variables were either moisture/aridity- or temperature-related, given that these two climatic factors appeared to be the main drivers of climate adaptation according to previous regional studies (Prunier et al., 2012; Hornoy et al., 2015).

**Table 1.** List of climatic factors extracted from BioSIM11 and used in this study.

| **Climatic variable** | **Abbreviation** | **Range (min; max)** |
| --- | --- | --- |
| Mean temperature during growing season<sup>1</sup> (°C) | Tmean_szn | 5.28; 16.39 |
| Yearly mean temperature (°C) | Tmean_yr | -6.94; 4.74 |
| Mean relative humidity during growing season (%) | RH_szn | 60.08; 81.04 |
| Yearly mean relative humidity (%) | RH_yr | 68.58; 79.84 |
| Mean climate moisture index during growing season | CMI_szn | 0.020; 4.23 |
| Mean drought code value during growing season | DC_szn | 89.07; 343.013 |
| Mean soil moisture index during growing season (%) | SMI_szn | 41.73; 98.13 |
| Mean vapor pressure deficit during growing season (hPa) | VPD_szn | 2.15; 7.68 |
| Mean vapor pressure deficit during peak of growing season<sup>2</sup> (hPa) | VPD_tg | 1.93; 8.93 |
| Mean vapor pressure deficit during hardening period (hPa) | VPD_ao | 2.13; 7.21 |
| Yearly mean vapor pressure deficit (hPa) | VPD_yr | 1.03; 4.08 |
| Number of thaw days during peak of winter<sup>3</sup> | TD | 0; 3.55 |
| Number of extreme frost days (temperature < -30°C) | Extr_FD | 1.58; 45.74 |
| Mean diurnal range (°C) | Bio2 | 7.99; 13.61 |
| Isothermality | Bio3 | 17.64; 31.50 |
| Temperature seasonality | Bio4 | 958.31; 1454.24 |
| Temperature annual range (°C) | Bio7 | 41.41; 54.71 |
| Precipitation seasonality | Bio15 | 36.07; 82.48 |
| Total precipitation during growing season (mm) | TP_szn | 112.75; 459.73 |
| Yearly total precipitation (mm) | TP_yr | 259.95; 1245.85 |
| Number of frost days during growing season | FD_szn | 4.61; 45.84 |
| Yearly number of frost days | FD_yr | 172.13; 263.45 |
| Total radiation during growing season (MJ/m<sup>2</sup>) | TR_szn | 2025.32; 2375.81 |
| Total radiation during peak of growing season (MJ/m<sup>2</sup>) | TR_tg | 1148.74; 1303.85 |
| Yearly mean aridity (mm) | Arid_yr | 12.00; 198.12 |
| Yearly total snowfall (mm of water equivalent) | TS_yr | 104.66; 434.88 |

**Table 1.**
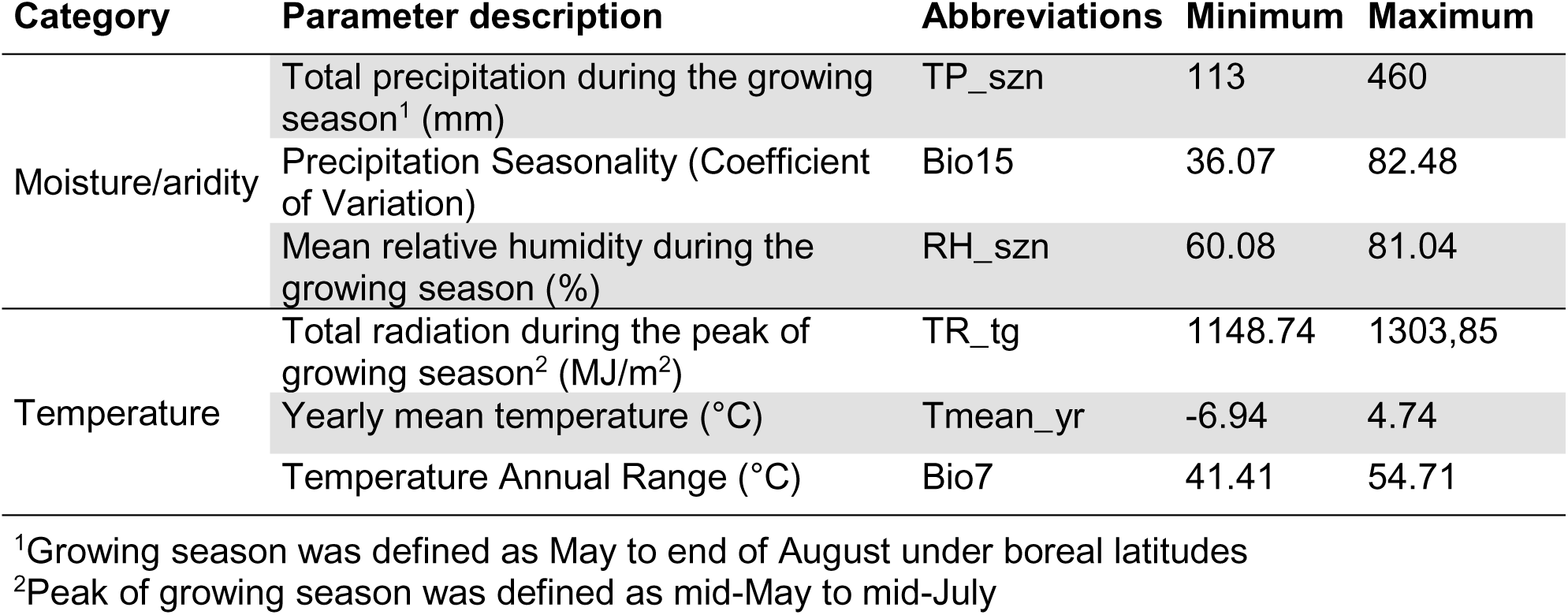
Retained climatic parameters and range of values across all sampled populations.

#### DNA extraction, genotyping assay and quality control

DNA was extracted from 100 mg of needles using the NucleoSpin 96 Plant II Extraction system (Macherey-Nagel; Duren, Germany) or the DNeasy 96 Plant Kit (Qiagen; Mississauga, Ontario, Canada). Concentration was confirmed by optical densitometry.

SNPs from 5183 distinct gene loci (one SNP per gene) were genotyped for each sample using the Illumina Infinium iSelect *PmGP1* genotyping array described in Pavy et al. (2016). A few advantages of using a validated gene SNP array over genotype-by-sequencing (GbS) in the context of this association study include assay repeatability, the fact that many gene loci are well annotated at the functional level, and a high genotyping rate which generally eliminates the need for data imputation (Pavy et al., 2008, 2013a, 2016; Lenz et al. 2020a,b). Following the protocol of Hornoy et al. (2015), gene SNPs that did not meet the following quality criteria were filtered out: minor allele frequency (MAF) ≥ 0.05, genotyping success rate (call rate) per SNP ≥ 90% and fixation index (*F*_IS_) ≤ |0.50|, indicative of potentially paralogous SNPs. Following these filtering criteria, 4838 high-quality SNPs remained in the genotyping dataset for subsequent analyses.

A few individuals with more than 10% of missing genotypes (call rate < 90%) were discarded with no data imputation attempted. Four populations that had only three individuals were also discarded because such a sample size was deemed insufficient to reliably estimate allele frequencies at the population level. In two instances, samples from the same population presented identical multi-locus genotypes; the sample that contained the most missing data was eliminated in both cases. After applying those filters, 254 individuals from 30 populations were retained (Fig. 1).

**Figure 1.**
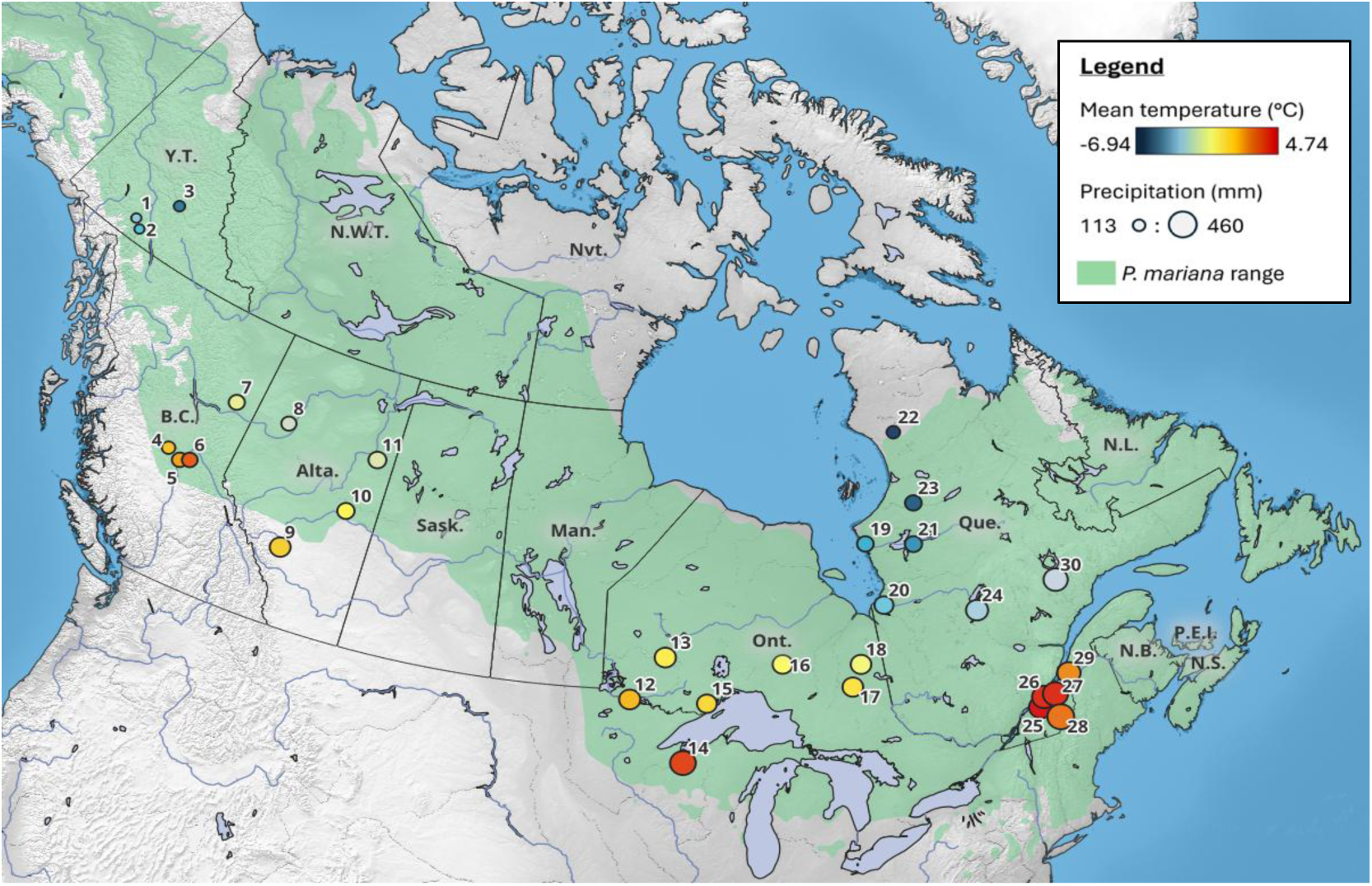
Geographical distribution of black spruce populations sampled Each circle represents a distinct natural population with number ordered by increasing longitude. As indicated in the insert in the top -right corner, circle size is representative of total precipitations during the growing season (larger circle means more precipitations) and circle color denotes mean annual temperatures (blue to yellow to red, blue representing colder temperatures). Black spruce natural range (Little, 1971) appears in green.

### Data analyses

#### Population structure

We used the Bayesian algorithm fastSTRUCTURE 1.0 (Raj et al., 2014) on the entire dataset of SNP genotypes, to infer historical population structure. After converting our dataset to STRUCTURE format, K (number of clusters) parameter values from 1 to 10 were tested using the chooseK function. This function is designed to identify the optimal K value by comparing the marginal likelihood obtained with different K values. Thus, the chooseK procedure helped determine the appropriate number of clusters that best represented the population structure in the dataset. The ancestry proportion coefficient (ancestry q-value) was also determined for each individual. It is defined as the estimated proportion of an individual’s genome that originates from each inferred population cluster.

#### Genotype-environment association (GEA)

We conducted genotype–environment association (GEA) analyses to explore how allelic frequencies correlate with different climatic variables, aiming to identify loci that exhibit signs of local adaptation. Missing data was addressed using the SNMF methodology and the impute function from the LEA package in R (Frichot & François, 2015) to fill in unknown genotypes.

We first used a partial Redundancy Analysis (pRDA), a multivariate method that aims to detect multilocus selection in large datasets by considering the covariation of groups of markers in response to environmental factors (Rellstab et al., 2015; Capblancq & Forester, 2021). Multilocus approaches are particularly suitable for studying adaptation in plants, as this process often results in subtle genetic changes across multiple loci rather than strong signals at individual loci (Savolainen et al., 2013; Hornoy et al., 2015). Additionally, most traits in conifers are inherited quantitatively (Neale & Wheeler, 2019), making this analysis especially adapted to our study.

Particular attention was paid to reducing correlation among climatic variables, as RDA is based on multiple linear regression and is therefore sensitive to strong collinearity among explanatory variable (Legendre & Legendre, 2012a,b). This was achieved by reducing the number of climatic factors through hierarchical clustering. First, all available climatic factors were standardized and their pairwise correlations were computed. The resulting correlation coefficients were then transformed into a distance matrix which was used to perform hierarchical clustering through the hclust function in R. Upon visual inspection of the resulting dendrogram, 11 clusters of correlated variables were determined (Fig. S1). One representative variable was selected from each cluster. Correlations among these preselected variables were then examined and any pairwise correlation exceeding a threshold of |0.7| resulted in the removal of one of the correlated variables. The retained climatic variables included three moisture/aridity- and three temperature-related indices (Table 2). Fig. 1 illustrates temperature and moisture/aridity gradients through proxy variables mean annual temperature and total precipitation during the growing season.

**Table 2.** Proportions of variance imputable to climate, geography, and genetic structure. Explainable variance is defined as constrained variance from the full model. Unexplained variance is the residual variance.

| Partial RDA models | Variance component | $R^2$ | $P(>F)$ | Proportion of variance | |
| --- | --- | --- | --- | --- | --- |
|  |  |  |  | Explainable | Total |
| Full model | 254.0 | 0.129 | 0.001*** | 1 | 0.15 |
| Climatic model | 56.5 | 0.029 | 0.001*** | 0.22 | 0.03 |
| Pure genetic structure model | 45.7 | 0.023 | 0.001*** | 0.18 | 0.03 |
| Pure geography model | 17.4 | 0.009 | 0.001*** | 0.07 | 0.01 |
| Confounded climate/structure/geography | 134.4 |  |  | 0.53 | 0.08 |
| Total unexplained | 1458.6 |  |  |  | 0.85 |
| Total variance | 1712.6 |  |  |  | 1 |

\*\*\* $P \leq 0.001$

Before conducting the RDA, we performed a variance partitioning analysis using the methodology from Capblancq & Forester (2021). The objective was to estimate the proportion of variance attributable to climatic factors, as opposed to potential confounding factors, namely neutral genetic structure and geography. We used four different RDAs to determine the proportion of variance explained by climatic factors, neutral genetic structure, and geography:

1. **Simple RDA — full model**: genotypes ∼ climate + lineage + geography
2. **Partial RDA — pure climate model**: genotypes ∼ climate, controlling for lineage + geography
3. **Partial RDA — pure genetic structure model**: genotypes ∼ lineage, controlling for climate + geography
4. **Partial RDA — pure geography model**: genotypes ∼ geography, controlling for climate + lineage

where “genotypes” are the SNPs’ genotype data, “climate” represents the six climatic factors, lineage is the ancestry coefficients from fastSTRUCTURE, and geography is the latitude and longitude of the studied natural populations.

Because the proportion of variance explained by the pure genetic structure model was considerable, we partialled out the effect of lineage in our RDA to avoid confounding effects arising from the historical genetic structure. Accounting for historical population structure allowed to assess adaptation in all populations at once, across historical lineages. The pRDA was conducted using the VEGAN package in R (Oksanen et al., 2009). Before computing, climatic variables were scaled and centered to a mean of 0 and a variance of 1.

After conducting the pRDA, the identification of SNPs under selection was achieved using the PCADAPT methodology (Capblancq et al., 2018), which relies on Mahalanobis distances. The four most significant RDA axes (*P* < 0.001 in permutation-based ANOVA with 999 permutations) were used to search for candidate SNPs. To account for multiple testing, q-values were calculated using the q-value package in R (Storey et al., 2024). Loci respecting a false discovery rate (FDR) threshold of q-value < 0.10 were considered significantly associated to climatic factors and compiled as candidate adaptive SNPs. Candidate SNPs were matched to the climatic predictor for which the correlation was the highest (as in Forester et al., 2018, see https://popgen.nescent.org/2018-03-27_RDA_GEA.html for details). If two factors showed comparable correlation levels (within a range of 10% of the best correlation), both factors were considered.

We also performed a univariate GEA method, namely Latent Factor Mixed Models (LFMM 2; Frichot et al., 2013; Caye et al., 2019). This algorithm uses least-squares minimization to estimate confounding (latent) factors, which allows controlling for population genetic structure. In order to capture most of the variance associated with the environment, a PCA on the scaled and centered 26 available climatic variables (Table 1) was first computed, and the scores of the first and second principal components were used as synthetic factors. The choice to retain the two first principal components was made by plotting the eigenvalues of principal components in a scree plot and using the broken stick method (Frontier, 1976). The first principal component explained 38.1% of the variance of the climatic factors and the second explained 33.8% of the variance. Upon examining the loadings on both axes as well as the PCA biplot (Fig. S2), it was determined that the first principal component represented mainly temperature variables and the second principal component represented mainly moisture/aridity-related variables.

We ran LFMM using the LFMM package in R (Caye et al., 2019) with the number of latent factors (K) set to two, as this was the optimal number of population structure clusters determined by fastSTRUCTURE. Similar to the pRDA, q-values were calculated, and a false discovery rate (FDR) threshold of 0.10 was applied to identify SNPs significantly associated with environmental variables. To identify the environmental predictor category most closely associated with each candidate SNP, we estimated the correlation between the genotypes and the scores of the first two principal components (with the first component representing temperature and the second one representing moisture/aridity as previously described).

#### F_ST_-based outlier detection method

To complement GEA analyses, an outlier detection method based on the fixation index (*F*_ST_) was implemented to identify loci under selection. This type of analysis compares the observed *F*_ST_ of a locus to its expected *F*_ST_ under a neutral model. Loci presenting significant divergence from the neutral model (outliers) are considered under selection. Because of the historical population structure present in our dataset, the hierarchical island model proposed in Excoffier et al. (2009) was deemed more adequate than other *F*_ST_-based methods. Indeed, this model accounts for more complex genetic structure, particularly when gene flow is impeded across different regions of a species’ range (i.e. when populations segregated into genetically divergent groups, as is the case in our study). The model operates on the premise that there is a greater exchange of migrants between demes within a group than among groups, to reduce the rate of false positives.

This method was implemented using the software ARLEQUIN 3.5 (Excoffier & Lischer, 2010). As prescribed in the hierarchical island methodology, populations were assigned to an eastern or a western group (matching historical population structure) based on their geographic location in order to mimic real migration paths. A total of 50 000 simulation iterations were run with 100 demes and 50 groups (as recommended in the ARLEQUIN 3.5 manual (Excoffier, 2011)). After running the algorithm, significant SNPs (*P* < 0.01) were compiled as candidate adaptive loci. Correlations between the genotypes of candidate SNPs and the scores of the first two principal components used in the LFMM analysis were computed to determine which predictor category is most closely associated with each candidate.

#### Intra-lineage analyses

In addition to the main analyses conducted on the full set of sampled individuals, we performed separate intra-lineage analyses for the western and eastern areas of the studied area. These analyses followed the same statistical procedures as the main analyses but assumed equal migration among populations within each lineage, using a regular island model instead of a hierarchical one (Excoffier et al., 2009; Excoffier, 2011). Given the relatively limited sample sizes within each lineage, these intra-lineage tests were only used to corroborate range-wide detections by verifying whether similar signals could be observed within specific portions of the range corresponding to historical lineages.

#### Functional annotations of candidate genes

All SNPs used in this study were located in coding regions of the genome, with each SNP representing a distinct gene locus. Given that LD generally does not exceed gene limits in conifer natural populations (including in black spruce; Pavy et al., 2012), each candidate SNP was associated with a distinct candidate gene assumed to be under selection. Functional annotations of the candidate genes were therefore investigated in order to corroborate their potential involvement in climate adaptation. To do so, we first retrieved black spruce contig sequences carrying candidate SNPs and searched their corresponding white spruce gene sequence (Rigault et al., 2011), as provided in Pavy et al. (2016). White spruce is a model organism within conifers and possesses some of the most comprehensive genetic resources available for the genus *Picea* (De La Torre et al., 2014; Gagalova et al., 2022; Pavy et al., 2025). Additionally, due to their longer sequence, white spruce gene clusters can be better annotated than fragmented contigs.

Using the sequences of the GCAT catalog genes (Rigault et al., 2011), BLASTX was run against the Uniprot-Swissprot protein database. Matching Uniprot IDs that satisfied an e-value < 1* 10^-10^ threshold were compiled. The ID mapping platform on the Uniprot website (https://www.uniprot.org/id-mapping) allowed the extraction of Gene Ontology (GO) terms for each retrieved ID, as well as information on protein families.

We used molecular function and biological process associated GO terms (Ashburner et al., 2000) to conduct an enrichment analysis using Fisher’s exact test. This method compares the frequency of occurrence of terms in an interest group (in our case, candidate genes under putative selection) compared to a background group (complete gene set). If the relative frequency of a certain term in the interest group was significantly higher than in the background group, it was declared overrepresented or “enriched”. Enrichment tests were conducted only on GO terms that appeared at least five times in the entire dataset.

For significant SNPs, the literature was screened for each underlying gene annotated with a Uniprot ID. Specifically, we examined the available evidence for their involvement in abiotic and biotic stress responses, as well as their possible roles in signaling pathways related to abscisic acid (ABA), jasmonic acid (JA), salicylic acid (SA) and auxin responses. We also investigated their contributions to plant development and growth, and their participation in critical biological functions such as photosynthesis, lipid metabolism, carbohydrate metabolism, transcription regulation, RNA processing and cell wall biogenesis. This examination primarily relied on studies on model organisms such as *Arabidopsis thaliana* and *Oryza sativa.* In addition, gene annotations and putative functions from previous studies on other conifer species including white spruce and Norway spruce (*Picea abies*) were consulted to support candidate gene function inference. This analysis was conducted for each candidate gene specifically, but if a role could not be retrieved for that exact protein, we further investigated whether this role was conducted by proteins of the same family.

## Results

### Population structure

The analysis of population structure performed using FastSTRUCTURE revealed an optimal number of two genetic clusters across the species’ distribution (Fig. 2), as this genetic partitioning generated the highest marginal likelihood among tested models. The partition of individuals in clusters matched almost perfectly the geographic distribution of populations across the studied area (see Fig. 1). Sorting the individuals based on their longitude of origin showed a clear separation between eastern and western populations (Fig. 2). Overall, 92 individuals belonged to the first cluster (in light blue on Fig. 2) whereas 161 individuals belonged to the second cluster (in dark blue on Fig. 2). We will subsequently refer to these clusters as the “western cluster” and the “eastern cluster”, respectively, due to their geographic location. Only one individual (Thunder Bay, Ontario) showed a significant degree of admixture. The western cluster ancestry proportion of this individual was 74% while it was geographically located in the eastern cluster. Such a result was considered anecdotal as all other individuals from the same population presented the expected eastern cluster signature. Ten individuals from both the western and eastern clusters also showed unexpected ancestry signatures (Fig. 2) in regard to known black spruce postglacial history (Jaramillo-Correa et al., 2004, 2009; Gérardi et al., 2010).

**Figure 2.**
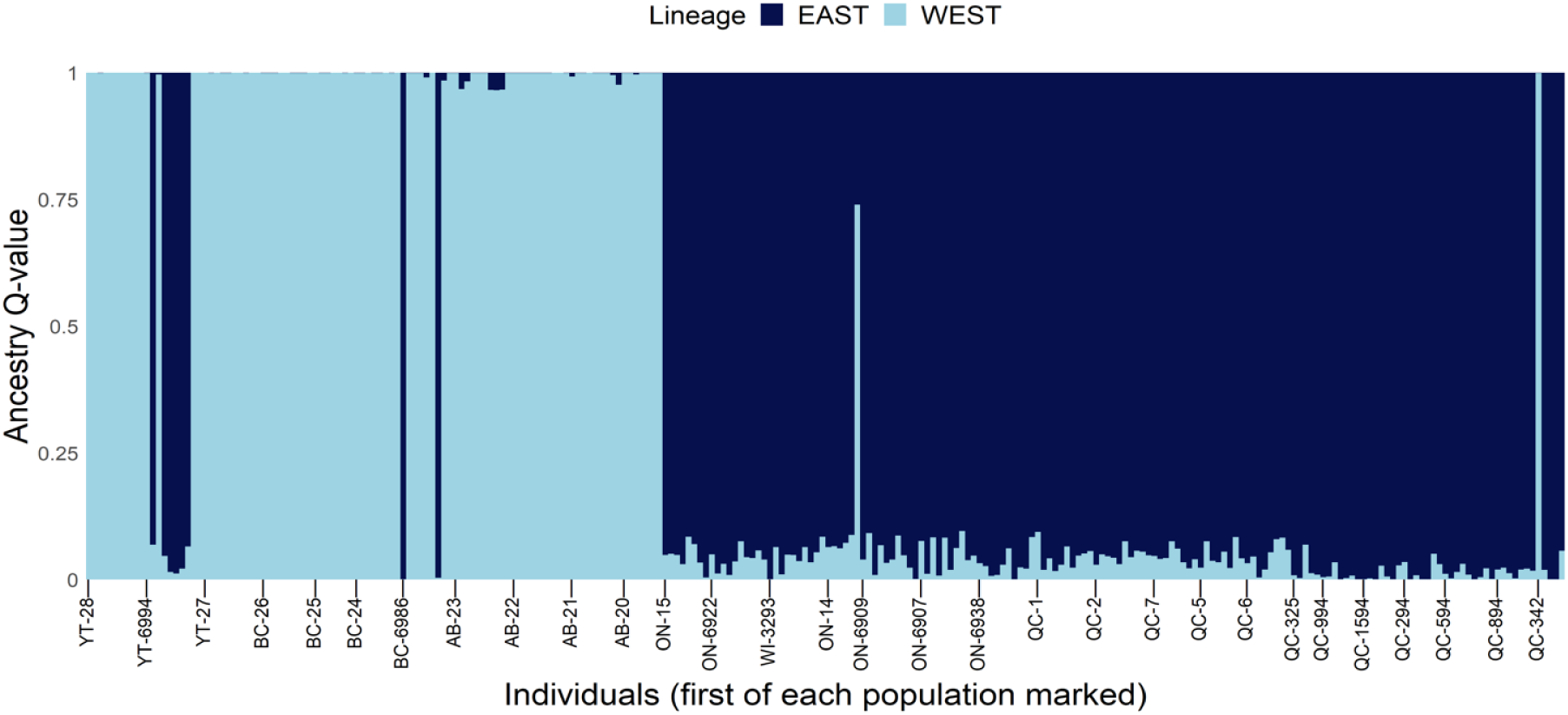
Population structure of sampled individuals. Ancestry coefficients (Q-values) estimated by FastSTRUCTURE. Individuals are ordered from westernmost to easternmost (left to right). Light blue represents the western cluster and dark blue represents the eastern cluster. The labels at the bottom are population codenames and are aligned with the first individual of each population.

This could be the result of isolated long-distance dispersal events, which are common in anemophilous tree species such as conifers (Gamache et al., 2003; O’Connell et al., 2007; Kremer et al., 2012) and can occur across thousands of kilometers (Comtois, 1997). In any case, association analyses conducted herein were designed to account for genetic structure, which should help mitigate the effects of such isolated discrepancies.

### Identification of candidate gene SNPs under selection

Before conducting GEA analyses, variance partitioning was used to estimate the proportion of variance attributable to climatic variables, as opposed to other potentially confounding factors, namely neutral genetic structure such as IBD, and geography. A full model incorporating climatic, genetic and geographic variables explained 15% of the overall genetic variation observed across sampled black spruce individuals (Table 3). Partial models for climate and genetic structure both explained 3% of total variance. They accounted for and 22% and 18% of the explainable variance, respectively. The geography partial model explained only 1% of total variance and around 7% of its explainable portion. The confounded effect of climate, genetic structure and geography represented 8% of the total variance and 53% of the explainable variance. These results highlight the considerable parallelism of environmental, genetic, and geographic gradients in our population sampling. In the subsequent RDA, genetic structure was incorporated as a covariate and partialled out to minimize the number of false positives. Geography was not included as a covariate because of its limited contribution to total variance and to minimize the risk of false negatives.

**Table 3.**
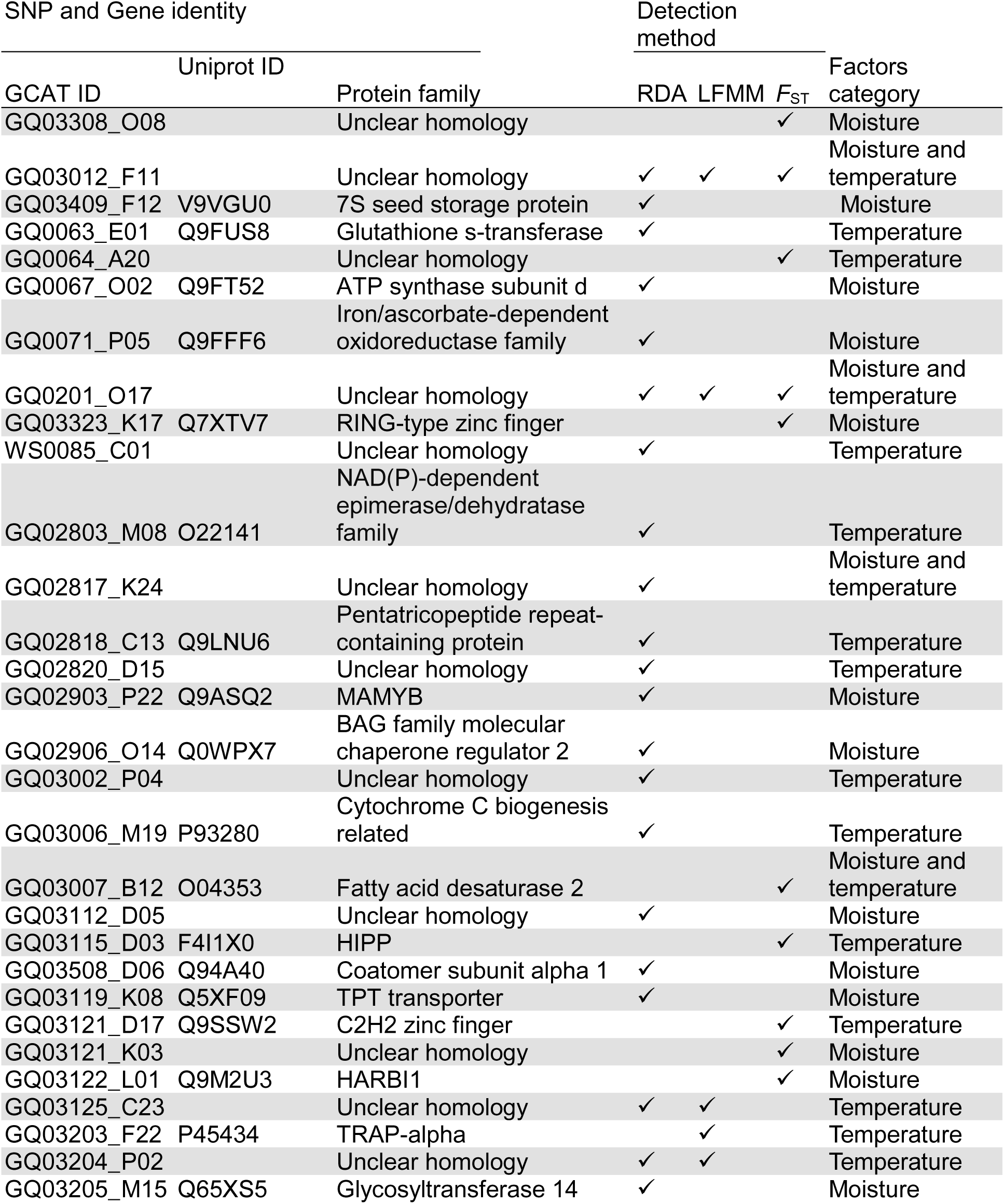
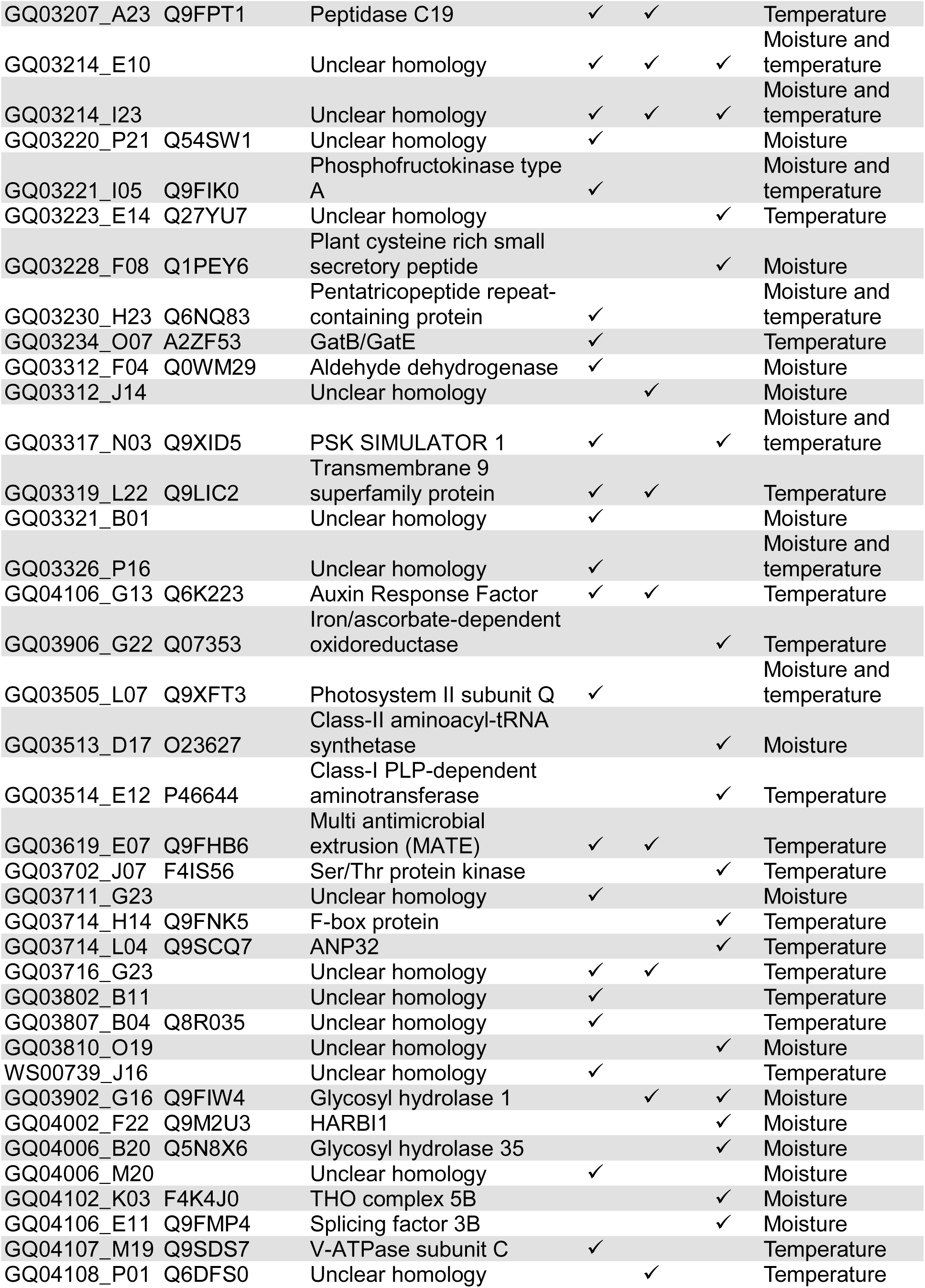
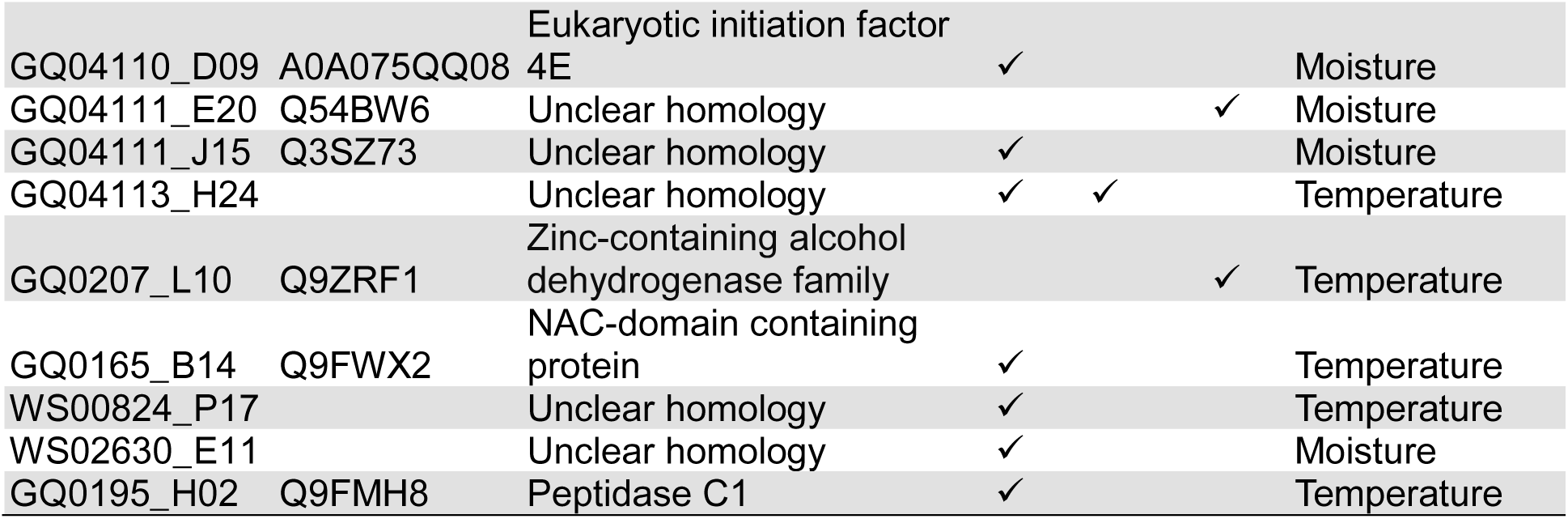
Functional annotations on the 77 candidate genes for climate adaptation in black spruce.

To identify gene SNPs potentially indicative of adaptation to climate, a pRDA was conducted on all sampled individuals with a False Discovery Rate (FDR) of 0.10. This resulted in the identification of 50 candidate gene SNPs, of which 23 were correlated with temperature variables (mean temperature, mean annual range and total radiation), 17 were correlated with moisture/aridity variables (total precipitation, relative humidity and precipitation seasonality), and 10 were correlated with both types of variables (Table 4).

With the same FDR threshold (Q < 0.10), the second GEA method LFMM detected 16 gene SNPs associated with synthetic factors (from PCA scores) representing the 26 climatic variables considered. Among these, 10 were correlated with temperature factors, two with moisture/aridity factors and four with both types of factors.

Regarding the *F*_ST_-based method, a total of 29 gene SNPs were detected as outliers at a significance threshold of *P* < 0.01. Computing correlations between the genotypes and the same synthetic variables from the LFMM method revealed that 13 SNPs were most correlated with precipitation factors, 10 SNPs were most closely correlated with temperature and six SNPs were correlated with both categories.

The gene SNPs detected by each method were compiled, which resulted in a list of 77 unique candidate SNPs, out of nearly 5000 gene SNPs tested, and covering as many distinct gene loci. This number represents about 1.5% success rate. Among those, four were detected through all three methods. There was also considerable overlap between the detections from pRDA and LFMM (12 candidates out of 16 LFMM candidates and 50 pRDA candidates), but considerably less between both GEA methods and the *F*_ST_-based method (five shared between the *F*_ST_-based method and pRDA, and five with LFMM). In total, 14 candidate gene SNPs were detected by at least two methods (Fig. 3). To reduce the number of false negatives, all gene SNPs detected were considered potential candidates for adaptation. However, SNPs cross-validated by multiple methods are likely to be more reliable and less prone to be false positives. In studies aimed at identifying potentially adaptive genes, such as this one, finding the right balance between minimizing false positives and avoiding the exclusion of true positives is challenging because there is no entirely objective way to achieve it.

**Figure 3.**
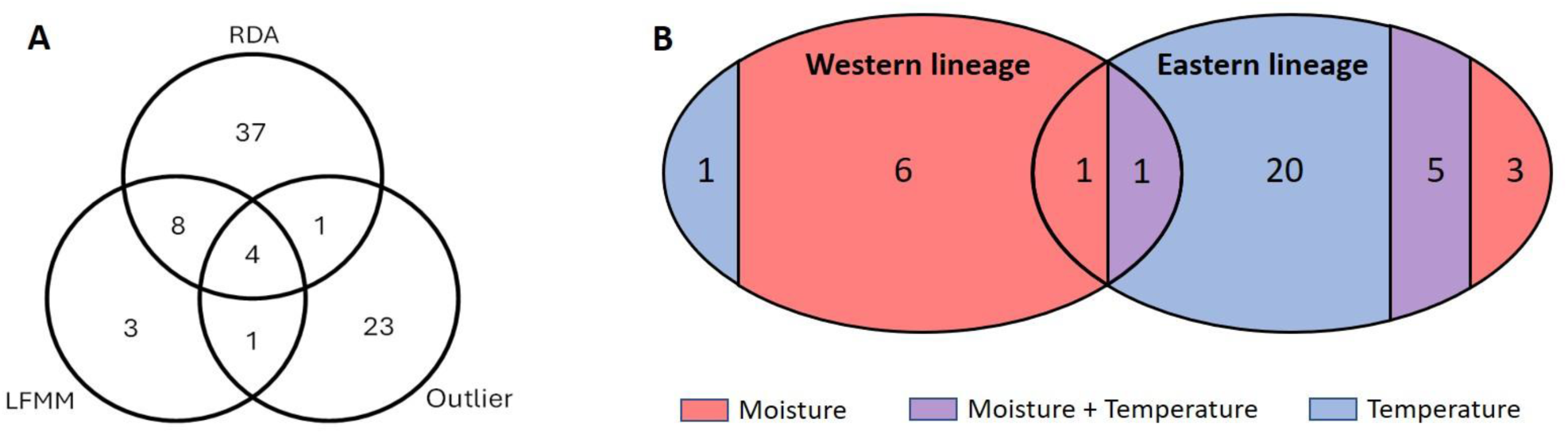
Overview of gene SNPs significantly associated with climatic variation in black spruce. (A) Out of the nearly 5000 gene SNPs from as many gene loci, distribution of the 77 significant gene SNPs identified from the joint rangewide analysis of both intra-specific lineages using three detection methods. (B) Lineage-specific corroboration of 37 gene SNPs initially identified in the rangewide analysis. Numbers indicate genes identified within each lineage or shared between lineages. Colors denote associations with moisture/aridity-related variables (red), temperature-related variables (blue), or both types of climatic factors at the same time (purple). The remaining 40 gene SNPs that were not corroborated by lineage-specific analyses comprised 21,14 and 5 genes associated with moisture/aridity, temperature and both types of climatic variables, respectively.

We repeated the aforementioned GEA (LFMM and RDA) and *F*_ST_ outlier detection methods within each of the western and eastern lineages of black spruce, thus without correction for the continental-wide population structure. These lineage-specific analyses were used as confirmatory analyses to ensure that the correction for population structure applied to the full dataset performed as intended. In total, 37 of the 77 candidate gene SNPs could be detected in at least one distinct lineage (Fig. 3b; Table S2), including 13 of the 14 candidate gene SNPs detected by more than one method across the whole species range. These analyses also showed that the eastern lineage harbored nearly four times as many corroborated candidate genes as the western lineage, likely due to the greater number of populations sampled and, consequently, higher statistical power in the eastern lineage. Of interest, the nature of associations between gene SNPs and climatic factors differed between lineages, with candidate genes from the eastern lineage predominantly linked to temperature, whereas those from the western lineage mostly associated with moisture/aridity, with an imbalanced ratio of 6:1.

### Gene annotations

The 77 gene SNPs detected in our analyses were all located in distinct black spruce genes, whose homolog sequences matched distinct genes from the white spruce gene catalog GCAT 3.3 (Rigault et al., 2011). The corresponding GCAT gene sequences of all gene loci carrying significant SNPs were blasted against the Uniprot-Swissprot database in order to retrieve functional annotations. Using the threshold e-value < 1×10^-10^, annotations were successfully obtained for 45 genes. These annotations are mainly from *Arabidopsis thaliana* and *Oryza sativa* but also from other conifers and vascular plant species. Homology to genes in other spruce species, including the largely distributed white spruce in North America and Norway spruce in Europe, was also assessed. Retrievable protein families for the candidate genes are listed in Table 4. Using ID mapping from the Uniprot website (https://www.uniprot.org/id-mapping), Gene Ontology (GO) terms were obtained for these genes. An enrichment analysis using these GO terms was performed using Fisher’s exact test. With a threshold *P* < 0.05, two molecular function GO terms were significantly enriched, namely copper-ion binding (GO:0005507) and nuclease activity (GO:0004518), and one biological process, mitochondrial translation (GO:0032543).

## Discussion

### Population structure and intraspecific lineages

Using ∼5000 SNPs distributed across as many distinct nuclear gene loci, we obtained high-resolution support for the major transcontinental population structure of black spruce previously inferred from mitochondrial, chloroplast, and a more limited set of nuclear DNA markers (Jaramillo-Correa et al., 2004; Gérardi et al., 2010; Prunier et al., 2012). Consistent with prior phylogeographic evidence, two genetically distinct and geographically structured population clusters were detected, matching black spruce western and eastern historical lineages, with a suture zone near the center of the North American continent in Canada. This concordance reinforces the well-documented west-east lineage split and confirms its persistence on a genomic scale in spite of gene flow. Notably, one western population (Marsh Lake, Yukon) contained genotypes more strongly associated with the eastern lineage than the western lineage (Fig. 2; YT-6994), as previously reported from mtDNA and nuclear SNP markers (Gérardi et al., 2010; Prunier et al., 2012).

The presence of an eastern cluster signature in a Yukon population could be explained by its proximity with the Beringian glacial refugium. Literature indicates that black spruce, as well as other plant and animal species, survived in a northern refugium located in Alaska during the Last Glacial Maximum (Abbott et al., 2000; Aubry et al., 2009; Gérardi et al., 2010; Napier et al., 2019). More precisely, it is hypothesized that black spruce had a continuous distribution between eastern and northwestern North America that got fragmented in isolated refugia during the last glacial period, including a Beringian refugium in the northwest of the continent. Preservation of a fraction of this ancestral black spruce population in this refugium would have allowed the conservation of its genetic background.

In accordance with this theory, studies on mtDNA, cpDNA and ncDNA detected variants typically associated with the central and eastern regions of the current species’ range in Alaska (Gérardi et al., 2010; Prunier et al., 2012). Furthermore, the contemporary eastern and western lineages are thought to derive from two distinct glacial refugia, presumably located south of the Great Lakes in the east and along the Pacific Coast or near the Cascade Range in the west (Jaramillo-Correa et al., 2004; Gérardi et al., 2010). The genetic pattern observed across Yukon populations in the present study is therefore consistent with the aforementioned theory and could be a result of gene flow from both the ancestral Alaskan population and the western lineage.

Interestingly, a single individual from the Manicouagan population in Quebec was assigned to the western lineage. Though anecdotal as only one individual from that population presented such genetic signature, this result is consistent with findings from Jaramillo-Correa et al. (2004), who reported the presence of a mitotype typically associated with the western black spruce lineage in northeastern Quebec and Labrador, not far from this population. Thus, this surprising pattern could result from admixture from the eastern lineage with a geographically more restricted lineage in Labrador (Jaramillo-Correa et al., 2004), in this northeastern region of North America.

### Identifying adaptive gene SNPs

We used multivariate and univariate GEA methods (pRDA and LFMM) as well as an *F*_ST_-based outlier detection method to identify SNPs presenting signatures of adaptation to climate. Combining these three methods provided a robust framework for identifying adaptive genetic variants. The value of using a combination of multivariate and univariate analyses has been highlighted in previous studies (Rellstab et al., 2015; Harrisson et al., 2017). Briefly, univariate methods are rather geared towards the detection of loci with strong individual effects on the phenotype under selection, whereas multivariate methods, by accounting for interactions among loci, allow to identify loci with smaller individual effects. Because the *F*_ST_-based method relies on population differentiation rather than explicitly modeling relationships between climatic variables and genotypes, it was used as a complementary approach to the two GEA analyses (RDA and LFMM). Given the exploratory nature of this study, all significant SNPs were considered as candidates for adaptation and retained for subsequent analyses, so to minimize false negatives in this discovery study. However, the 14 SNPs identified by more than one method should be considered as higher-confidence candidate loci involved in adaptation to climate. In total, the total of 77 significant SNPs detected in this study represented only 1.5% of the SNPs tested. This is a very low rate compared to most other conifer and spruce studies using similar GWAS approaches (for a review, see Table 5 in Bousquet et al., 2021), thus suggesting a low rate of false-positives in the present study.

While both GEA methods (RDA and LFMM) showed a substantial overlap in adaptive gene SNP detections (12 candidates out of a total of 16 for LFMM and 50 for RDA), the overlap between the *F*_ST_-based method and the GEA methods was more limited (5 shared candidates with RDA and 5 with LFMM out of 29 total *F*_ST_ candidates). As stated before, LFMM and RDA are both based on linear regressions whereas *F*_ST_-based methods rely on differentiation among populations. This difference means that GEA methods are designed to detect adaptation along environmental gradients (clines), whereas *F*_ST_-based methods can detect strong signals of local adaptation within single or a few populations. As such, GEA methods are well-suited to detect small and gradual shifts in allele frequencies along an environmental gradient (Hornoy et al., 2015), but would potentially not detect adaptation signals from a restricted number of populations being significantly differentiated. Furthermore, *F*_ST_-based methods may identify SNPs as false-positives when their spatial distribution is driven by processes unrelated to climate, such as biotic interactions or other untested environmental factors, or genetic processes related to demographic history such as acute genetic drift (Hornoy et al., 2015), even if the hierarchical *F*_ST_ method used herein (Excoffier et al., 2009) should limit the number of such false-positives. Nevertheless, these fundamental differences between detection methods could potentially explain, at least in part, the dissimilarities observed in the overlap among identified subsets of candidate gene SNPs.

Out of 77 significant gene SNPs, a total of 31 were correlated with moisture/aridity factors (42 when considering significant relationships with both moisture/aridity and temperature factors). The detection of moisture/aridity-related candidate genes was expected on account of both environmental data and black spruce biology. Indeed, moisture/aridity-related variables follow a quite marked longitudinal gradient across the boreal Canadian landscape, with the western populations occurring in much drier conditions than their eastern counterparts (Prunier et al., 2012). For instance, during the growing season, the mean precipitation west of the exclusion range of the provinces of Saskatchewan and Manitoba was only 214 mm, whereas it was 342 mm in the eastern part of the range. While resilient to a large spectrum of environmental conditions, previous studies have indicated that black spruce could be sensitive to acute drought stress (Balducci et al., 2013; Way et al., 2013; Desaulniers et al., 2026). This potential vulnerability of black spruce to water stress could originate from its shallow root system (Viereck & Johnston, 1990), which could prevent it from drawing water deeper in the soil. Additionally, Peng et al. (2011) noted that the mortality of boreal trees (including black spruce) across Canada was on the rise from 1963 to 2008, and suggested water stress from regional drought episodes as a dominant contributor to this phenomenon. Such conclusions were also recently reinforced (Marchand et al., 2025). Evolutionary pressures towards adaptive traits associated to water-related stress are thus likely to be important in black spruce, as evidenced for the sympatric white spruce where genetic variation and local adaptation to drought was found significant (Depardieu et al., 2020, 2024). Hence, the combined effects of regional differences in moisture availability and the sensitivity of black spruce to water stress likely drive local genetic adaptation, as previously observed in a more geographically restricted dataset in eastern Canada (Desaulniers et al., 2026), highlighting the need for detailed analyses of quantitative traits related to drought resilience across the species’ natural range.

On the other hand, 35 candidate gene SNPs were most correlated with temperature-associated factors (46 when considering detections with both moisture/aridity and temperature factors). In a similar pattern to moisture/aridity-related variables, temperature exhibited a pronounced gradient across the spatially extensive studied area, though it was latitudinal instead of longitudinal. Thus, northern populations of black spruce generally experience considerably colder conditions than those in the south, especially in the eastern part of the range (Fig. 1). In spite of its wide ecological amplitude, black spruce can be sensitive to heat stress, causing damage that can affect wood properties or anatomy, especially in the southern part of its range (Colombo & Timmer, 1992; Mirabel et al., 2022; Reed-Métayer et al., 2025). As for other spruces, it can also be sensitive to cold stress and frost damage at the seedling and more mature stages (Dang et al., 1992; Man et al., 2021), including the sensitivity of late frost events after budbreak, which can damage severely spruce shoots and meristems, and sometimes lead to mortality in younger spruce individuals (Mura et al., 2022; see also Benomar et al., 2022). Other studies (e.g. Beaulieu et al., 2004; Chagnon et al. 2023) showed that growth in black spruce is strongly influenced by latitudinal temperature gradients, reinforcing the view that temperature-driven selection pressures intensify with latitude. Taken together, these findings suggest that the extensive latitudinal range and associated temperature gradient across the studied area likely contributed to local adaptation in black spruce, which may explain the numerous temperature-associated candidate gene SNPs detected in our analyses.

In addition, 11 gene SNPs were correlated with both moisture/aridity- and temperature-related climatic factors. Such identification of gene SNPs related to both factors is consistent with findings from previous studies in black spruce (Prunier et al., 2011, 2012) and in white spruce (Hornoy et al., 2015). Overall, responses to dehydration and cold in plants is often tightly interconnected, for instance at the level of transcriptional processes (Yamaguchi-Shinozaki & Shinozaki, 2006).

Drought resistance has also been shown to be correlated with cold tolerance in Norway spruce (Blödner et al., 2005), further reinforcing the idea that genes simultaneously involved in adaptation to both factors are plausible.

Intra-lineage regional analyses conducted without rangewide correction for population structure helped assessing the repeatability of candidate gene SNP detections and if climate factors driver of adaptive evolution existed between the two large regions. By repeating GEA and *F*_ST_ independently in the eastern and the western parts of the studied area, we thus evaluated whether signals identified in the full dataset persisted within lineages. The fact that 37 out of 77 candidate gene SNPs (and 13 out of 14 high-confidence gene SNPs) could be detected in at least one of the two regions suggests that our procedure for controlling population structure-induced variance was effective. However, the number of corroborated candidate genes was uneven between lineages, with almost four times more genes recovered in the eastern lineage than in the western lineage. While this asymmetry may truly reflect stronger or more abundant lineage-specific genotype-environment associations in the eastern part of the range, it was possibly also influenced by the larger sampling size in the eastern lineage, which likely translated into greater statistical power to detect significant associations in this region.

However, beyond this imbalance, the results also suggest that the predominant climatic driver of adaptation differed between the two lineages (Fig. 3B). In the western lineage, corroborated candidate gene SNPs were predominantly associated with moisture/aridity-related variables, outnumbering significant temperature-related gene SNPs by a ratio of roughly 6 to 1. Conversely, the eastern lineage displayed the opposite pattern, with a clear predominance of significant gene SNPs associated with temperature-related variables in a reverse manner, by a ratio of approximately 6 to 1 compared to those associated to moisture/aridity factors. A complementary explanation could also be that the two lineages were exposed to climatic gradients of different amplitude, which may influence both the intensity of selection and the power to detect associated gene SNPs.

In addition, Figure 3B highlights that only two significant gene SNPs were shared between the western and eastern lineage-specific analyses, suggesting limited parallel evolution of the same adaptive loci in both lineages. This pattern is consistent with the hypothesis that lineage-specific adaptive SNPs may emerge when adaptation is fueled by genetically distinct pools of standing genetic variation among lineages, each of which reflecting at least partly its demographic history (Prunier et al., 2012). Notwithstanding differences in statistical power, the relatively large fraction of significant gene SNPs identified only in the range-wide analysis suggests that some associations were detectable only across larger climatic gradients, which should increase the ability to identify adaptive signals that were weak or relatively uniform within individual lineages. Altogether, these results show the importance of integrating both lineage-specific and cross-lineage approaches when working with historically structured populations spread on large geographical areas.

Across spruce species, numerous studies identified SNPs and genes associated with climatic factors, most prominently temperature and moisture/aridity parameters. In black spruce, GEA studies have identified a range of adaptive loci, including nonlinear relationships (Prunier et al., 2011, 2012). Similar climate-associated variants have been detected in white spruce using both outlier and GEA approaches (Namroud et al., 2008; Hornoy et al., 2015; Depardieu et al., 2021). Likewise, in Norway spruce, numerous SNPs have been identified, including stress-response loci (Azaiez et al., 2018) and clinal variants associated with latitude gradients and phenology (Chen et al., 2012). The genotyping array used in this study was developed based on exome capture probes designed on white spruce (Stival Sena et al., 2018). Thus, most black spruce genes genotyped in this study have white spruce homologs, and some also have homologs in Norway spruce as well (Azaiez et al., 2018). Some overlap between our candidate genes and those reported in these earlier studies was observed, particularly at the gene family level. These shared observations with past studies are explored further in the following section of the discussion.

### Putative functions and roles of candidate genes carrying significant SNPs

Functional annotations could be retrieved for 45 of the 77 genes found to carry significant SNPs in relation to local climate variation. The literature on model plants and related conifers was screened to infer their putative biological roles. A great variety of molecular functions and biological processes was observed among these candidate genes (Fig. 4), several of them being of particular interest in the context of molecular adaptation to climate. This is consistent with findings in congeneric species such as the sympatric transcontinental white spruce, where a similar diversity in putative adaptive genes has been reported (e.g., Hornoy et al., 2015). Moreover, given the very high molecular genetic diversity of black spruce and, as in other conifers, the presence of large gene families and functional redundancy (Pavy et al., 2013, 2025; Lo et al., 2024), a broad diversity of candidate genes significantly associated with climatic variation was expected. Notably, these genes quite differ from those previously identified in congeneric species, even though the latter were also associated with diverse biological functions, as observed in other conifers (Pavy et al., 2025).

The GCAT gene GQ0063_E01 codes for protein glutathione S-transferase U17 (GSTU17). Glutathione s-transferases are proteins induced by diverse environmental stimuli in order to maintain cell redox homeostasis and protect organisms against oxidative stress (Chen et al., 2012). GSTU17 specifically has been linked to the development of seedling in *A. thaliana* (H.-W. Jiang et al., 2010). A knockout mutant of this gene also improves resistance to salinity and drought stresses, as well as higher levels of abscisic acid (ABA) and glutathione. This suggests the involvement of GSTU17 in modulating ABA-mediated stress response in plants. Pavy et al. (2025) also identified in conifers a positively selected gene homologous to a glutathione s-transferase in *A. thaliana*. These findings also suggest a conserved function in modulating stress tolerance across different plant groups.

The gene GQ03121_D17 was identified as a zinc finger protein AZF2. This protein is a transcription factor of the zinc finger C2H2 family that acts as an expression repressor of specific genes under stress conditions (Kodaira et al., 2011). More specifically, AZF2 downregulates the expression of ABA-repressive and auxin-inducible genes. Furthermore, AZF2 transcription was shown to be induced by abiotic stresses such as drought, cold, and high salt (Sakamoto et al., 2004). This suggests a role in both ABA and auxin signaling pathways in addition to involvement in abiotic stress response. Other than that, zinc finger C2H2 proteins (Namroud et al., 2008; Prunier et al., 2011, 2012, 2013), transcription factors from the MYB (Namroud et al., 2008; Prunier et al., 2011, 2012, 2013; Lo et al., 2024; Pavy et al., 2025), NAC, and C3HC4 RING (Namroud et al., 2008; Prunier et al., 2011, 2012, 2013) families have also been identified as potentially adaptive genes in black spruce and white spruce from previous studies. Candidate genes identified in this study that belong to these families were GQ03323_K17, GQ02903_P22 and GQ0165_B14, respectively. Notably, RING-type zinc finger proteins (associated with the GQ03323_K17 gene) have been reported as involved in climate adaptation in white spruce (Hornoy et al., 2015), consistent with its well-established roles in plant growth, development, and responses to abiotic stress (Han et al., 2022).

The gene GQ03702_J07 represents integrin-linked protein kinase 1 (ILK1). This protein is part of the Rapidly Accelerated Fibrosarcoma-like mitogen-activated protein kinase (RAF-like MAPKKK) family. This protein family is involved in a cascade that amplifies signals from receptors to targets (Caffrey et al., 1999). In plants, it has been linked to ABA and auxin signaling pathways (Kuhn et al., 2024; Takahashi et al., 2020). Studies on ILK1 mutants in *A. thaliana* linked it to increased osmotic stress sensitivity, resistance to bacterial pathogens as well as growth (Brauer et al., 2016; Chinchilla et al., 2008).

Further supporting the relevance of the detected adaptive genes in black spruce, homologs of three significant genes from the present study were also identified in an earlier GEA analysis conducted by Depardieu et al. (2021) in white spruce. These genes are GQ03514_E12, a Class-I PLP-dependent aminotransferase, GQ02903_P22, a MAMYB transcription factor, and GQ03308_O08, which remains unannotated. The recurrence of detection of these candidate genes in independent studies and among two phylogenetically distant spruce species suggests an implication in parallel environmental responses in conifers.

An enrichment analysis using Fisher’s exact test was performed using the GO terms extracted by ID mapping. Statistical power was limited due to the small number of GO terms represented five times or more in the background dataset, which was a criterion that we applied to reduce the occurrence of false positives. Nevertheless, two molecular function terms, and one biological process term were found enriched in our dataset.

Molecular function significantly enriched terms were copper-ion binding (GO:0005507) and nuclease activity (GO:0004518). Proteins with the copper ion binding molecular function interact non-covalently with copper. Copper acts as a cofactor for numerous enzymes (e.g. Cu/Zn-superoxide dismutase (Cu/ZnSOD), cytochrome-c oxidase, ascorbate oxidase, amino-oxidase, laccase, plastocyanin, and polyphenol oxidase (Yruela, 2009) and in key biological processes (e.g. photosynthetic and mitochondrial electron transport, oxidative stress responses, hormone perception, and cell wall metabolism (Himelblau & Amasino, 2000; Pilon et al., 2006; Weigel et al., 2003), which can be involved in stress responses. Interestingly, Azaiez et al. (2018) found this GO term to be enriched among SNP-rich genes in Norway spruce, indicating that these genes are generally characterized by high standing variation. The enrichment of this term in black spruce significant genes could imply that this variation is caused by environmental selection. Regarding nuclease activity, it has been associated in plants with DNA repair and leaf senescence (W. Sakamoto & Takami, 2014; Spampinato, 2017), which are both important in stress response (Guo et al., 2021; Spampinato, 2017).

The biological process found to be statistically enriched was mitochondrial translation (GO:0032543). Studies in various plant species have demonstrated alterations in the abundance of mitochondrial proteins when oxidative stress is induced (e.g. Hossain et al., 2012; Sweetlove et al., 2002; Taylor et al., 2009). Hence, this could indicate that this biological process is truly involved in response to stress in black spruce.

Several of the 45 annotated genes identified as harboring significant adaptive SNPs presented functional annotations that were either not clearly characterized, or too broad (Fig. 4 - Other functions), making it difficult to determine whether these genes could play a significant role in stress response and at what level. These annotations include transport proteins, histone deacetylases, tRNA biosynthesis-related proteins, among others. As they were detected as genes carrying significant SNPs in our analyses, there is a possibility that these genes (as well as the unannotated ones) are involved in unexplored stress response pathways related to climatic pressures in conifers and other long-lived trees, which are underrepresented in the plant literature.

Our results are also in agreement with those of Pavy et al. (2013b, 2025), Gagalova et al. (2022), and Lo et al. (2024), who observed that a large proportion of the rapidly evolving genes under positive selection were related to stress response in different spruce genomes and transcriptomes, including in black spruce. While deciphering the same general pattern as we observed in the present ecogenomic study, these gene sequence studies used a drastically different analytical approach based on detecting sequence signatures of positive selection from patterns of nonsynonymous and synonymous substitutions significantly diverging from expectations under neutral evolution.

**Figure 1.1.**
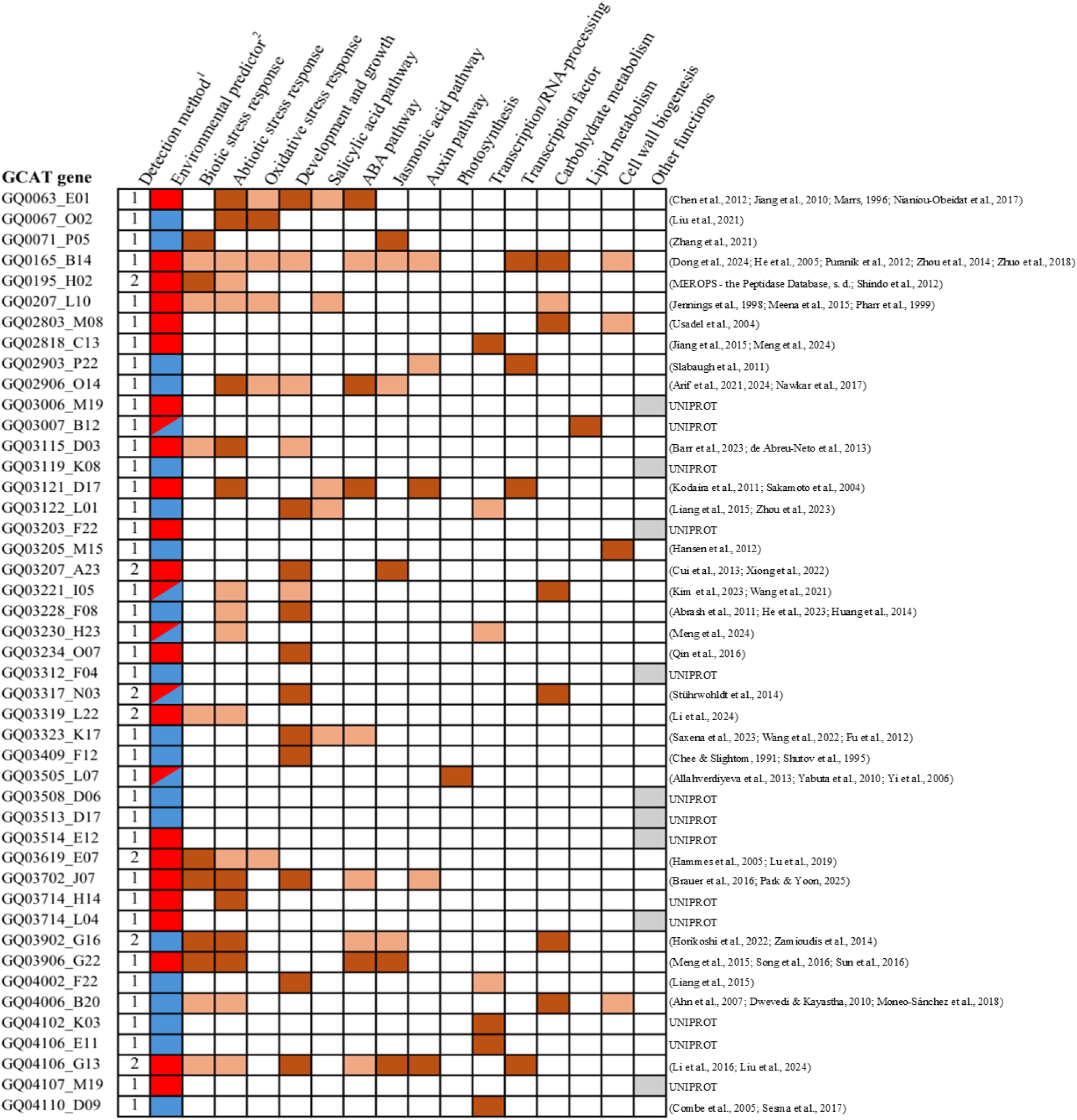
Functional annotations of 45 identified black spruce genes associated with climate adaptation. Each line represents a distinct gene and columns represent molecular functions or physiological roles. ^1^This column represents the number of detection methods that identified each candidate gene. ^2^Color codes for categories of climatic factors; blue means correlation with moisture/aridity, red means correlation with temperature, and bicolor cells indicate relationships with both types of climatic factors. ^3^Remaining columns: a dark red cell indicates that there is evidence in literature supporting the involvement of the specific gene in this function/role; a lighter orange cell means that evidence was found only at the gene family level. References are on the right, UNIPROT means no mention of interest was found for this gene other than in the Uniprot-Swissprot database.

### Study limitations and research perspectives

Although historical population structure was explicitly accounted for in our analyses, the presence of genetically distinct lineages among sampled populations might have impacted our results, as the historical population structure overlaps with the well-known east-west decreasing precipitation gradient across the species’ range, as the climate becomes more continental. While methods correcting for population structure were applied to reduce spurious associations, they may have also limited the detection of true associations, especially for moisture/aridity-related factors. Indeed, because intraspecific lineage differentiation parallels the longitudinal precipitation gradient across Canada, a stringent approach and correction were necessary. At the same time, overcorrection may have reduced the number of moisture/aridity-related candidate SNPs and increased the likelihood of false negatives. Additionally, sample size was somewhat limited due to the exclusion of samples from the contact zone between the eastern and western lineages, as well as the zone of contact between black spruce and red spruce in southern Québec and the Maritime provinces. While this reduced sample size may have limited statistical power and likely lead to the detection of fewer significant gene SNPs, excluding these zones of the range likely increased the confidence that the detections reflected true signals of adaptation rather than introgression or historical population structure.

The genes significantly associated to climatic factors identified herein open interesting new avenues for future research. For example, the functional role of some genes carrying significant SNPs remains unknown. Subsequent transcriptomic studies, such as differential gene expression analyses (Stival Sena et al., 2018; Depardieu et al., 2021; Laoué et al., 2021; Ribeyre et al., 2025) or GPA studies (e.g. Depardieu et al., 2021) could corroborate the adaptive role of these candidate genes in response to environmental pressures.

This study may also provide valuable insights for spruce breeding programs. For instance, the identified SNPs associated to climatic factors could, in theory, be used in Marker Assisted Selection (MAS); a technique where genetic markers associated with specific phenotypic or environmental traits are used to select individuals carrying the desired traits to improve adaptation (Collard & Mackill, 2008). However, in trees, traits related to climate adaptation are typically under polygenic control, with each locus explaining only a small fraction of phenotypic variation (Muranty et al., 2014). This would limit the operational efficiency of MAS in this context.

Alternatively, the adaptive candidate gene SNPs identified could be used to improve the accuracy of genomic selection (GS) models, which have been already developed and tested positively in black spruce (Lenz et al., 2017; Soro et al., 2026). GS uses genome-wide marker data to predict the genetic value of individuals in breeding populations and it has been applied in several spruce species for various traits (Beaulieu et al. 2014a,b; Lenz et al., 2017, 2020a,b; Bousquet et al., 2021; Soro et al., 2026) and in many other conifers and angiosperm trees (for a review, see Beaulieu et al. 2024), including for improving resilience to insect pests (Beaulieu et al., 2020; Lenz et al. 2020a) and climate anomalies such as severe drought episodes (Laverdière et al., 2022). GS does not require prior knowledge of the genetic basis of the trait of interest, as it uses statistical models to predict the cumulative effect of all SNPs on a phenotype (Beaulieu et al., 2014a, 2024).

Interestingly, a previous study in black spruce showed that over-representing markers with larger effects could improve GS model accuracy (Lenz et al., 2017). Also, preselecting markers significantly associated with desirable traits has been shown to improve GS model accuracy (Hiraoka et al., 2018; Z.-Q. Chen et al., 2023), similarly as for the use of candidate gene SNPs for increasing the success of GPA analyses (Prunier et al., 2013).

Such strategies are likely to increase the contribution of short-range LD between markers and quantitative trait loci (QTL) in GS modeling, rather than relying solely on long-range LD and relatedness (Beaulieu et al., 2014b; Lenz et al., 2017; Soro et al., 2026). Thus, while marker-assisted selection (MAS) is not readily operational for polygenic quantitative traits where single QTL and SNP effects remain usually small in spruces (Beaulieu et al., 2011; Pelgas et al. 2011; Pavy et al., 2017), the candidate gene SNPs identified herein could still add value by improving the accuracy of GS models, and in turn, better predicting genetic values associated to black spruce resilience to climate variation, for instance. This would represent a practical way to leverage adaptive genetic information related to climate in marker-assisted breeding strategies, which we intend to test in black spruce using information gathered from the present study.

Future works could also benefit from the candidate gene SNP subset identified herein to expand landscape genomics studies and for use in genetic conservation efforts. Indeed, GEAs can be extended to estimate genomic offset, defined as the expected mismatch between a population’s current genetic composition and its forecasted environmental conditions. Recent applications of such models allow researchers to project spatial patterns of adaptive potential under different climate change scenarios (Fitzpatrick & Keller, 2015), thereby identifying populations at higher risk of maladaptation, notably within the genus *Picea* (Lachmuth et al., 2023). Thus, genomic offset could be used to pinpoint vulnerable populations in the face of climate change but also offers a powerful tool to guide assisted gene flow by translocating genotypes adapted to future conditions into these populations.

## Conclusion

This study has enhanced our understanding of the genomic basis of climate adaptation in black spruce, a keystone species of North America’s boreal forests. Through the application of an informed genotyping approach and robust statistical methods, we have identified a set of candidate genes significantly associated with key climatic factors related to temperature, drought and cold stresses, uncovering multiple processes and metabolic pathways involved in response to local climates. In addition to advancing our fundamental knowledge on climate adaptation in black spruce, these results also have practical implications for genetic conservation and breeding programs, for instance by potentially improving genomic prediction models in the context of increasing climate instability and occurrence of severe climate anomalies (IPCC, 2023). Thus, this work should help anticipate the evolutionary trajectory of this largely distributed boreal tree species under climate change, and provide marker information for conservationists and breeders to assist them in selecting tree stock with improved adaptive potential to future local climates.

## Acknowledgements

We would like to thank Prof. Juan Pablo Jaramillo-Correa (Autonomous Univ. of Mexico) and Dr. Julien Prunier (CRCHUL, Univ. Laval) for their helpful comments and suggestions on a previous draft of this manuscript. We also thank France Gagnon (Univ. Laval) for her help with genotyping samples and data validation. This work was part of the FastTRAC II spruce genomics project lead by J.B. and P.L. and funded by Genome Canada and Genome Quebec (Genomic Applications Partnership Program - GAPP). We also acknowledge support from the Canada Research Chair in Forest Genomics to J.B., the Natural Sciences and Engineering Council of Canada (NSERC) for a discovery grant to J.B., and the Fonds de recherche du Québec - Nature & Technologies (FRQ-NT) for a competitive graduate research scholarship to V.Q.

## Supplementary display items

**Table S1.**
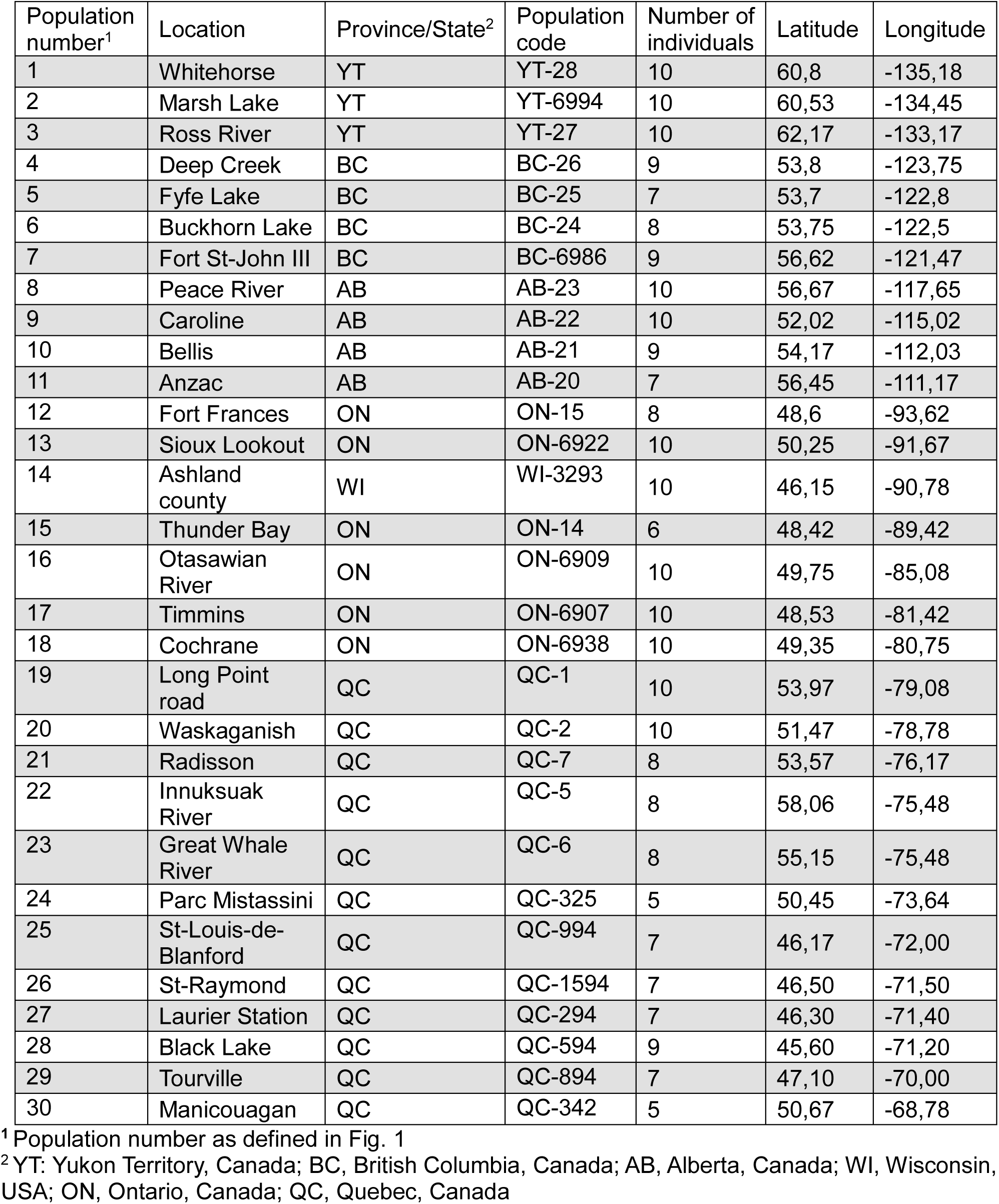
List of black spruce natural populations used in this study.

**Table S2.** List of black spruce candidate genes resulting from inter-lineage analysis and corroborated by at least one intra-lineage analysis.

| GCAT ID | RDA | LFMM | $F_{ST}$ |
| --- | --- | --- | --- |
| GQ0063_E01 | ✓ |  |  |
| GQ0165_B14 | ✓ |  |  |
| GQ0195_H02 | ✓ |  |  |
| GQ0207_L10 |  |  | ✓ |
| GQ02818_C13 | ✓ |  |  |
| GQ03006_M19 | ✓ |  |  |
| GQ03203_F22 |  | ✓ |  |
| GQ03207_A23 | ✓ | ✓ |  |
| GQ03221_I05 | ✓ |  |  |
| GQ03234_O07 | ✓ |  |  |
| GQ03317_N03 | ✓ |  | ✓ |
| GQ03319_L22 | ✓ | ✓ |  |
| GQ03619_E07 | ✓ | ✓ |  |
| GQ03807_B04 | ✓ |  |  |
| GQ03902_G16 |  | ✓ | ✓ |
| GQ04107_M19 | ✓ |  |  |
| GQ04108_P01 |  | ✓ |  |
| GQ04110_D09 | ✓ |  |  |
| GQ03214_E10 | ✓ | ✓ | ✓ |
| GQ03716_G23 | ✓ | ✓ |  |
| GQ04113_H24 | ✓ | ✓ |  |
| GQ03125_C23 | ✓ | ✓ |  |
| GQ03204_P02 | ✓ | ✓ |  |
| GQ03012_F11 | ✓ | ✓ | ✓ |
| GQ03214_I23 | ✓ | ✓ | ✓ |
| GQ0201_O17 | ✓ | ✓ | ✓ |
| GQ03802_B11 | ✓ |  |  |
| GQ03112_D05 | ✓ |  |  |
| GQ04006_M20 | ✓ |  |  |
| WS00824_P17 | ✓ |  |  |
| GQ0067_O02 | ✓ |  |  |
| GQ03205_M15 | ✓ |  |  |
| GQ03312_F04 | ✓ |  |  |
| GQ04006_B20 |  |  | ✓ |
| GQ03002_P04 | ✓ |  |  |
| WS02630_E11 | ✓ |  |  |
| GQ03308_O08 |  |  | ✓ |

**Figure S1.**
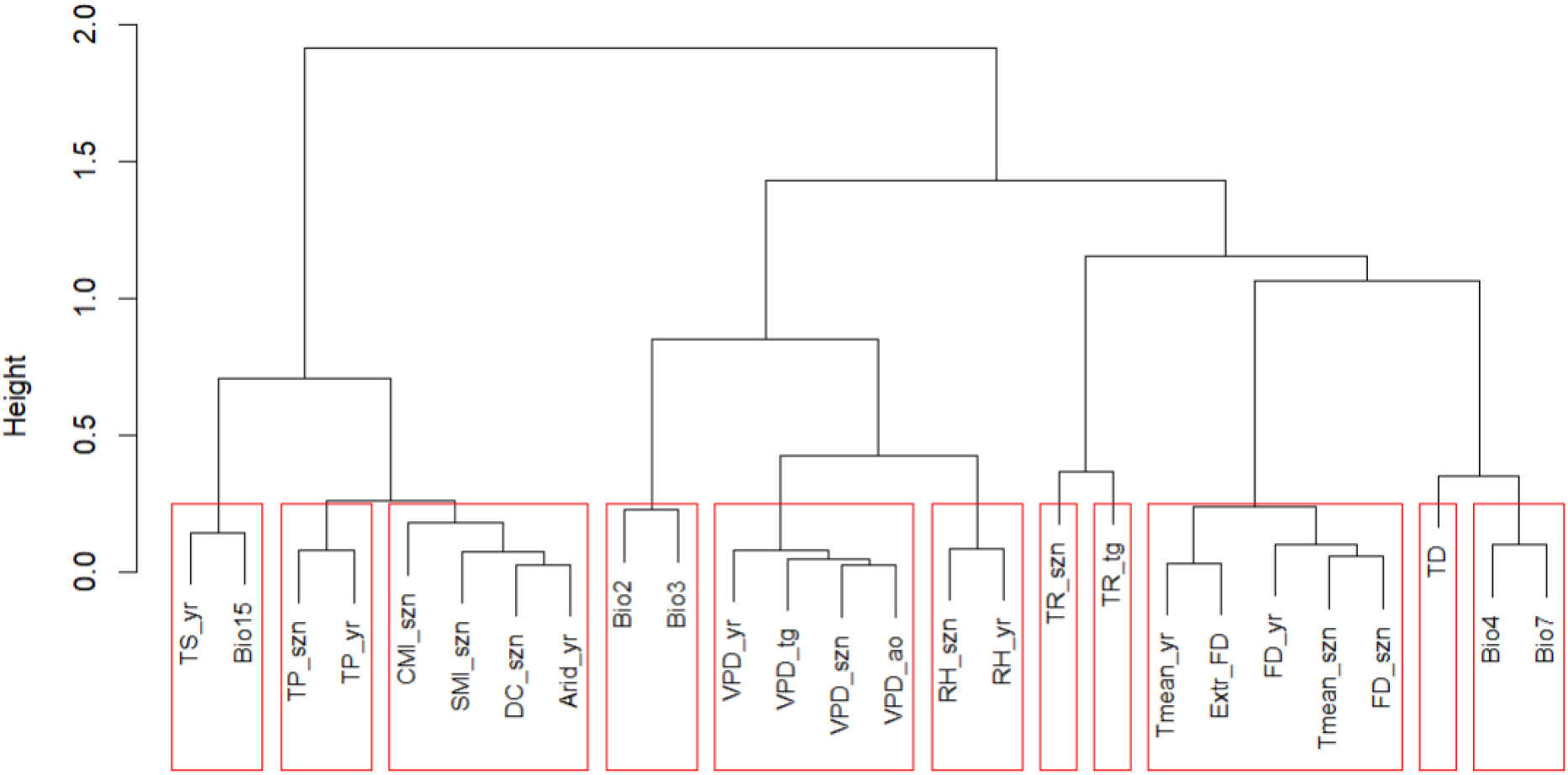
Correlation-based clustering of climatic parameters used for RDA parameter selection. Red rectangles indicate variable groupings. Definitions of climatic factor abbreviations are provided in Table 1 of the main text.

**Figure S1.**
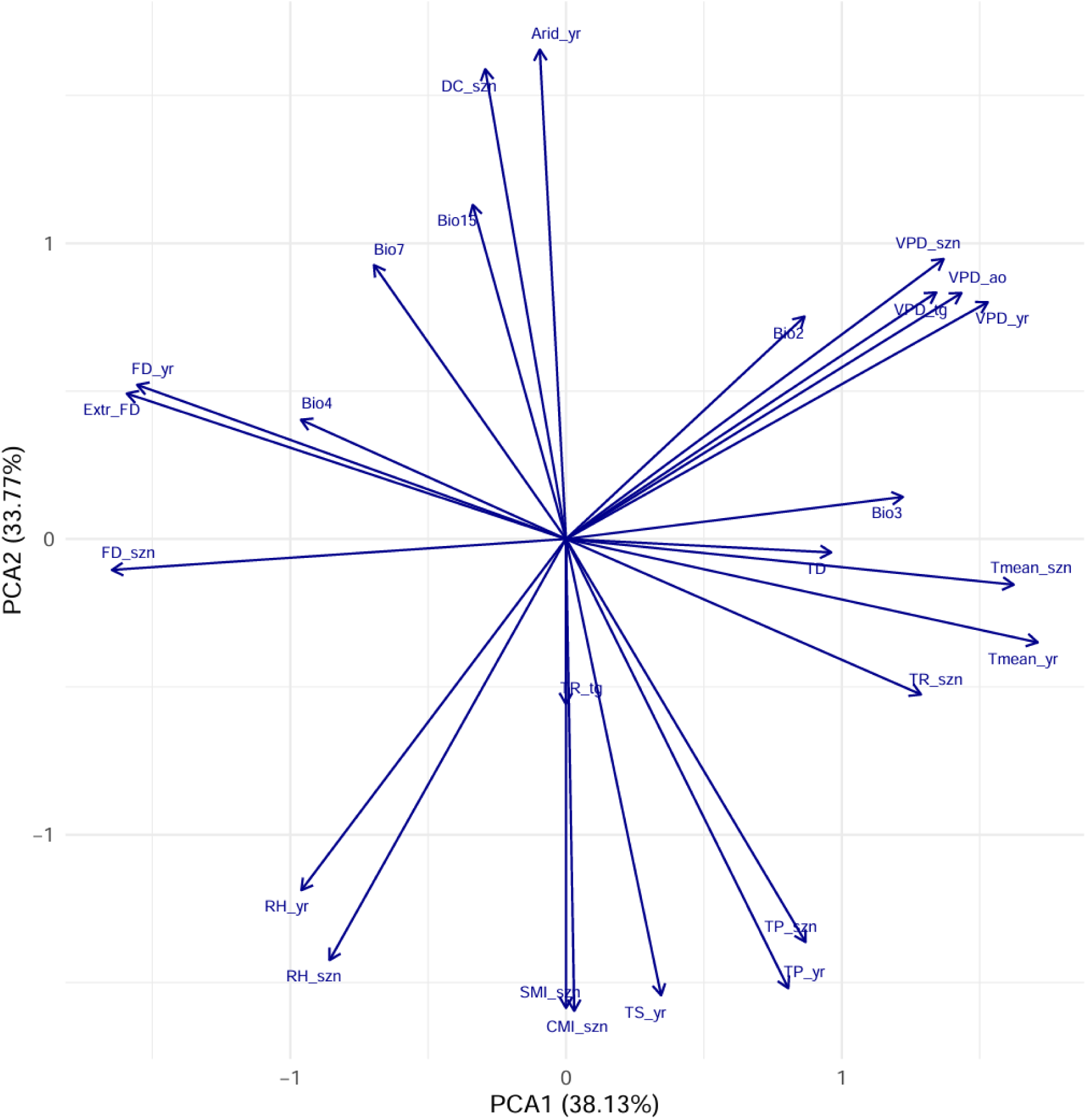
Principal component analysis of the 26 climatic variables used in this study. Definitions of climatic factor abbreviations are provided in Table 1 of the main text.

